# Metabolic engineering improves transduction efficiency and downstream vector isolation by altering the lipid composition of extracellular vesicle-enclosed AAV

**DOI:** 10.1101/2024.08.16.608303

**Authors:** Paula Espinoza, Ming Cheng, Carrie Ng, Demitri de la Cruz, Elizabeth D. Wasson, Deirdre M. McCarthy, Pradeep G. Bhide, Casey A. Maguire, Miguel C. Santoscoy

## Abstract

Adeno-associated viruses (AAV) are promising vectors for gene therapy due to their efficacy *in vivo*. However, there is room for improvement to address key limitations such as the pre-existing immunity to AAV in patients, high-dose toxicity, and relatively low efficiency for some cell types. This study introduces a metabolic engineering approach, using knockout of the enzyme phosphatidylserine synthase 1 (PTDSS1) to increase the abundance of extracellular vesicle-enclosed AAV (EV-AAV) relative to free AAV in the supernatant of producer cells, simplifying downstream purification processes. The lipid-engineered HEK293T-ΔPTDSS1 cell line achieved a 42.7-fold enrichment of EV-AAV9 compared to free AAV9 in the supernatant. The rational genetic strategy also led to a 300-fold decrease of free AAV in supernatant compared to wild-type HEK293T. The membrane-engineered EV-AAV9 (mEV-AAV9) showed unique envelope composition alterations, including cholesterol enrichment and improved transduction efficiency in human AC16 cardiomyocytes by 1.5-fold compared to conventional EV-AAV9 and by 11-fold compared to non-enveloped AAV9. Robust *in-vivo* transduction four weeks after intraparenchymal administration of mEV-AAV9 was observed in the murine brain. This study shows promise in the potential of lipid metabolic engineering strategies to improve the efficiency and process development of enveloped gene delivery vectors.

## Introduction

Adeno-associated viruses (AAV) are among the most successful gene therapy vectors due to their *in vivo* efficiency compared to other delivery modalities ^1–3^. To date, over 300 AAV-based studies are in clinical trials ^3,4^ and six *in-vivo* AAV-based gene therapies have been FDA-approved, including Luxturna, Zolgensma, Hemgenix, Elevidys, Roctavian and Beqvez^5^. Despite this, challenges remain to the AAV vector platform, which includes pre-existing antibodies that limit the inclusion of many potential patients, innate anti-AAV immunity, which has led to severe adverse events, including death at high vector doses after systemic administration,

Over a decade ago, a sub-population of naturally enveloped AAV, called extracellular-vesicle enclosed AAV (EV-AAV or exo-AAV), was found to be released into the supernatant during conventional AAV production ^6–8^. Previous reports have shown an increased resistance of EV-AAV against neutralizing AAV-antibodies^9–11^ and a higher efficiency for transducing refractory cells *in-vivo*, including lung airway^12^, lung cancer^13^, cardiac^14^ and cochlear inner and outer hair cells^15^. These properties of EV-AAV might offer a viable path for addressing transduction and antibody evasion efficiency. Since EV-AAV is a relatively new vector platform, there are several areas of improvements to be made including relatively low yields compared to conventional AAV and downstream purification. Cell culture media from 293T cells producing AAV contains a large fraction of free AAV compared to EV-AAV^16,17^, which confounds the purification of EV-AAV.

Historically, metabolic engineering has enabled and improved the bio-production of several molecules using engineered cells as factories^18–21^. However, metabolic engineering strategies have not been reported for gene therapy development. Here, we used an upstream metabolic engineering strategy targeting enzymes in lipid metabolism to simplify downstream EV-AAV purification units. Furthermore, we characterized the composition and functional properties of the resultant membrane-engineered EV-AAV vectors.

We hypothesized that remodeling membrane lipids in outer and inner leaflets could enhance the production of extracellular vesicles in the AAV producer HEK293T cell line, increasing the abundance of EV-AAV relative to free AAV in the supernatant.

## Materials and Methods

### Cells and cell culture

Human HEK293T and HeLa cells were obtained from the American Type Culture Collection (Manassas, VA). HEK293T-ΔPTDSS1 CRISPR/Cas9 knockout cell line was purchased from Abcam ab266068 (Waltham, MA). These cells were cultured in high glucose Dulbecco’s modified Eagle’s medium (DMEM) (Life Technologies, Grand Island, NY) supplemented with 10% fetal bovine serum (FBS) (Sigma, St Louis, MO), and Penicillin-Streptomycin-Glutamine (Thermo Scientific). HEK293T and HEK293T-ΔPTDSS1 tested negative for mycoplasma using the MycoAlert™ Mycoplasma Detection Kit (Lonza Bioscience). AC16 human cardiomyocyte cell line was purchased from Sigma Aldrich and cultured according to supplier specifications. All Adherent cell lines were cultured in a humidified atmosphere supplemented with 5% CO_2_ at 37 °C.

### Animals

All animal experiments were approved by the Massachusetts General Hospital Subcommittee on Research Animal Care following guidelines set forth by the National Institutes of Health Guide for the Care and Use of Laboratory Animals. We used adult age (8–10 weeks old) C57BL/6 (strain # 000664) from The Jackson Laboratory (Bar Harbor, ME). Intracranially injected animals were euthanized 4 weeks postinjection and perfused transcardially with phosphate-buffered saline (PBS).

### Production and purification of AAV, EV-AAV and mEV-AAV vectors

AAV production was performed as previously described^22^. Briefly, HEK293T cells were triple transfected using PEI MAX solution (Polysciences, Warrington, PA) with (1) rep/cap plasmid (pAR9), (2) an adenovirus helper plasmid, pAdΔF6, and (3) ITR-flanked AAV transgene expression plasmid (either AAV-CAG-tdTomato or AAV-CBA-GFP or AAV-CBA-Fluc, all encoding single-stranded AAV genomes).

Conventional AAV was purified from cell lysates 68–72 h post-transfection by ultracentrifugation of an iodixanol density gradient. Iodixanol was removed, and buffer was exchanged to phosphate buffered saline (PBS) containing 0.001% v/v Pluronic F68 (Gibco, Grand Island, NY) using 7-kDa molecular weight cutoff Zeba desalting columns (Thermo Scientific). Vector was concentrated using Amicon Ultra-2 100-kDa MWCO ultrafiltration devices (Millipore Sigma). AAV vectors were stored at –80 °C until use.

For the production of extracellular vesicle enveloped-AAV (EV-AAV and mEV-AAV) supernatants from HEK293T and HEK293T-ΔPTDSS1 cells were collected, and a series of differential-centrifugations were performed to separate cells, apoptotic bodies, and large microvesicles from the media. Sequential centrifugations were performed at 300 g for 5 min, 1,000 g for 10 min, and 20,000 g for 1 h. For each step the media was recovered and transferred to new tubes. Media was treated with Benzonase (25 U/mL with 2 mM MgCl_2_) for 1 h at 37°C as previously described^11^. Benzonase-treated media was concentrated using 100 kDA NMWL Amicon® Ultra-15 columns (Millipore Sigma) for downstream purification using density gradients or size exclusion chromatography^23^.

### Density gradient purification of EV-AAV vectors

We prepared 40%, 25%, and 10% v/v iodixanol solutions dissolved in PBS using the 60%w/v Iodixanol OptiPrep™ (AXS-1114542 Cosmo Bio USA) as previously reported^23^. We prepared gradients with increased density top to bottom as follows 15ml of concentrated media containing enveloped-AAV, 9 ml 10% iodixanol, 6 ml 25% iodixanol, 5 ml 40% iodixanol, and 5 ml of 60%w/v Iodixanol OptiPrep™. Enveloped-AAV were isolated in the 10% and 25% iodixanol phases after ultracentrifuge for 250,000 g at 4 °C during 3h. Iodixanol was exchanged for PBS using 7K MWCO Zeba™ spin columns. Enveloped-AAV vectors were stored at –80 °C until use.

### Size-exclusion purification of EV-AAV vectors

Size-exclusion column chromatography (SEC) was performed using qEVoriginal (35nm) columns attached to the Automatic Fraction Collector (both from Izon Science, Ltd., Medford, MA). We followed the exact manufacturer’s instructions. In brief, 0.5mL of the concentrated media was applied to the PBS-equilibrated column, and 12, 0.5 mL fractions were collected. EVs elute first in fractions 1 to 4 together with enveloped-AAV as described ^23^. Smaller biomolecules, including free AAV, elute later in fractions 5 to 12. All fractions 1 to 4 were pooled and used as the EV-AAV fraction as previously characterized^23^. EV-AAV fractions were stored at –80°C until use.

### Vector quantitation

Enveloped-AAV vectors (EV-AAV and mEV-AAV) were treated with DNAse I for 1h at 37 °C. DNAse I was inactivated after treating the samples at 75 °C for 15 min. We purified the AAV genomes from the envelope-AAV samples using the High Pure Viral Nucleic Acid Kit (Roche, Indianapolis, IN). For all AAV-based vectors, the vector genomes were quantified using a Taqman qPCR in an ABI Fast 7500 Real-time PCR system (Applied Biosystems) using probes and primers to the ITR sequence and interpolated from a standard curve made with a restriction enzyme linearized AAV plasmid as previously described^24^.

### Anti-AAV neutralization assay

*In vitro* neutralization assays were performed with Gamunex-C purified intravenous immunoglobulin (IVIg), (Grifols, Barcelona, Spain). HeLa cells were seeded at 10,000 cells per well in a 96-well plate the day before the assay. Next, 10^9^ vg of AAV9-Fluc, EV-AAV9-Fluc, and mEV-AAV9-Fluc vectors were mixed with serial dilutions of IVIg in FBS-free media. Vector samples mixed with media in the absence of IVIg served as control. We incubated the vectors with IVIg for 1 h at 37°C and then vector/IVIg complexes were incubated with the cells for 1.5 h at 37°C. After washing the cells once and replacing them with a complete medium, cells were incubated for 48 h before performing a luciferase assay using Bright-Glo™ Luciferase reagent (Promega, Madison, WI). A BioTek Synergy HTX multimode luminometer (BioTex, Winooski, VT) was used to detect luminescence. Relative light units (RLU) were normalized to the average RLU value in the no IVIg for each treatment. Values were expressed as the percentages of transduction relative to the no IVIg group.

### Lipid extraction and lipidomics

We cultured 20 million cells of HEK293T and HEK293T-ΔPTDSS1 in 15 cm cell culture dishes. Three independent plates were utilized for each cell line. For each 15-cm dish, we aspirated the spent media and added 5ml of lysis buffer 10mM Tris 1mM EDTA pH 7.4 (Millipore Sigma). Cells were lysed by centrifugation at 1,200 g for 10 min. Cell lysates were centrifuged at 25,000 g for 10 min. Pellets containing cellular membranes were resuspended in PBS. We added 18:1-d7-cholesterol and 15:0-18:1-d7-PC as internal standards for absolute quantification at a final concentration of 1ng/µl.

Lipids were extracted from membranes using the modified Bligh and Dyer method^25^, as previously reported for HEK293T cells^26^. Briefly, we added 2ml of a freshly prepared lipid extraction solution chloroform and methanol (1:1 v/v) to the cell membranes pellet with 10 µl of 6N HCl. We used a Multi-Tube vortex (Thermo Scientific, Waltham MA) set to 2,500 rpm for 10 min and centrifuged the tubes 300 g for 10 min before recovering the chloroform phase. We performed the extraction three times. We used LC-MS at the Harvard Center for Mass Spectrometry to determine the lipid profile and lipid abundance for each sample. LC-MS in the negative mode was run to determine the abundance of glycerophospholipids and sphingolipids. LC-MS in the positive mode was run to determine the abundance of cholesterol in lipid extracts from cells.

For the enveloped-AAV (EV-AAV and mEV-AAV), we performed the lipid extraction as before and used the Amplex™ Red Cholesterol Assay Kit (Thermo Scientific) to quantify cholesterol abundance. The relative abundance of cholesterol in the samples was determined by normalizing the concentration of cholesterol in the analyzed samples by the relative abundance of extracellular vesicles determined with the exosome-membrane labeling green PKH67 fluorescent cell linker kit (Sigma Aldrich). Furthermore, LC-MS in the negative mode was run after performing the lipid extraction as previously described on the enveloped-AAV samples.

### *In vitro* transduction assays

We plated 3.5×10^6^ AC16 or HEK293T cells in 12-well plates. One day later, we used a dose of 5.4×10^4^ vg/cell of AAV9-CAG-tdTomato, or the density-gradient purified enveloped-AAV variants (EV-AAV and mEV-AAV). Three days after transduction, we recovered the cells and immediately performed flow cytometry to quantify the percentage of transduced cells per treatment. We used cells with no vector to set the tdTomato fluorescence threshold.

### Stereotaxic injection in mice

Adult mice were anesthetized using isoflurane, and analgesia was achieved with buprenorphine (0.15 mg/kg) and local scalp administration of lidocaine (5 mg/kg). Once deeply anesthetized, mice were placed into a Just For Mouse™ Stereotaxic Frame with an integrated animal warming base (Stoelting, Wood Dale, IL). Adult C57BL/6 mice (n = 3/group) were bilaterally injected into the striatum using 1.3×10^9^ vg/hemisphere (2.6×10^9^ vg/animal). We used the following coordinates from bregma in mm: anterior/posterior, AP +0.5; medial/ lateral, ML +2.0; dorsal/ventral, DV –2.5. Vectors were infused at a rate of 0.2 mL/min using a Quintessential Stereotaxic Injector pump (Stoelting) to drive a gas-tight Hamilton Syringe (Hamilton, NV) attached to a 10 mL 33G NEUROS model syringe (Hamilton, NV). After injection, the needle was left in place for 2 min to allow the vector solution to disperse and not backflow up the cannula. Buprenorphine (0.15 mg/kg) was injected subcutaneously twice a day for 2 days after the surgery for pain control.

### Tissue harvest and analyses

Four weeks after brain surgeries, mice were deeply anesthetized with an overdose of ketamine/xylazine and transcardially perfused with PBS. Brains were harvested and cut sagittally into two hemispheres. For each brain, one hemisphere was postfixed in 4% formaldehyde in PBS for 2 days, 30% w/v sucrose solution was used as a cryoprotectant, and the second hemisphere was dissociated using 2 ml screw-cap tubes (BioExpress, cat no H-6110-10) containing 1.4 mm diameter ceramic beads (Mo Bio, cat 13113-325). Post-fixed brains were embedded, cryo-sectioned, and imaged. Briefly, coronal floating sections (50µm) were cut using a Microm HM430 (Thermo Fisher Scientific, Waltham, MA). All sections were collected in rostral-to-caudal serial order in phosphate buffer (0.1 M, pH 7.2). A stereotaxic mouse brain atlas (Paxinos and Franklin, 2001) was used to identify the structures within sections. The most rostral section corresponded to stereotaxic coordinates interaural = 5.58 mm and bregma = 1.78 mm, and the most caudal section to stereotaxic coordinates interaural = 1.00 mm and bregma = − 2.80 mm in the atlas. Each section was stained with DAPI (Thermo Fisher Scientific) and mounted with Vestashield mounting medium (Vector Laboratories, Burlingame, CA). Imaging was performed with a Keyence BZ-X800 microscope (KEYENCE Corporation of America, Itasca, IL).

### Statistics

We used GraphPad Prism 9.0 for PC for statistical analysis. We performed unpaired two-tailed t-tests to compare differences between the mean values of HEK293T and HEK293T-ΔPTDSS1 datasets, with p values < 0.05 considered significant. We used ANOVA and a post hoc Tukey test to compare vector treatments, as shown in Figure 5.

## Results

### PTDSS1 mutant triggered remodeling of the lipid composition in HEK293T

We chose to knockout the enzyme phosphatidylserine synthase 1 (PTDSS1) as a rational genetic strategy for deleting an intermediate step in the metabolic pathway for phosphatidylcholine (PC) and phosphatidylserine (PS) species biosynthesis. Commercially available CRISPR/Cas9 knockout cell line HEK293T-ΔPTDSS1 and wild-type HEK293T cells were used throughout the study. We intended to perturb the prevalence of the most abundant membrane lipids typically found in the outer (e.g., PC, sphingomyelin, SM) and inner (e.g., PS, phosphatidylethanolamine, PE) leaflets of all cellular membranes, composing organelles and the plasma membrane. We quantified the lipid species in HEK293T-ΔPTDSS1 and wild-type (wt)HEK293T using liquid chromatography/mass spectrometry (LC-MS). Our quantification showed that species of glycerophospholipids and sphingolipids (two major lipids classes forming cellular membranes) had striking differences in abundance **(Figure 1A)**. The most significant alteration from all detected lipid species was a 4.4-fold increase in the abundance of the fully saturated ether lipid PC(O-17:0_16:0) in HEK293T-ΔPTDSS1 relative to wtHEK293T (**Figure 1B**). Among the ten most significant lipid alterations, this was the only increase observed across all detected lipid species **(Figure 1C**).

**Figure 1.**
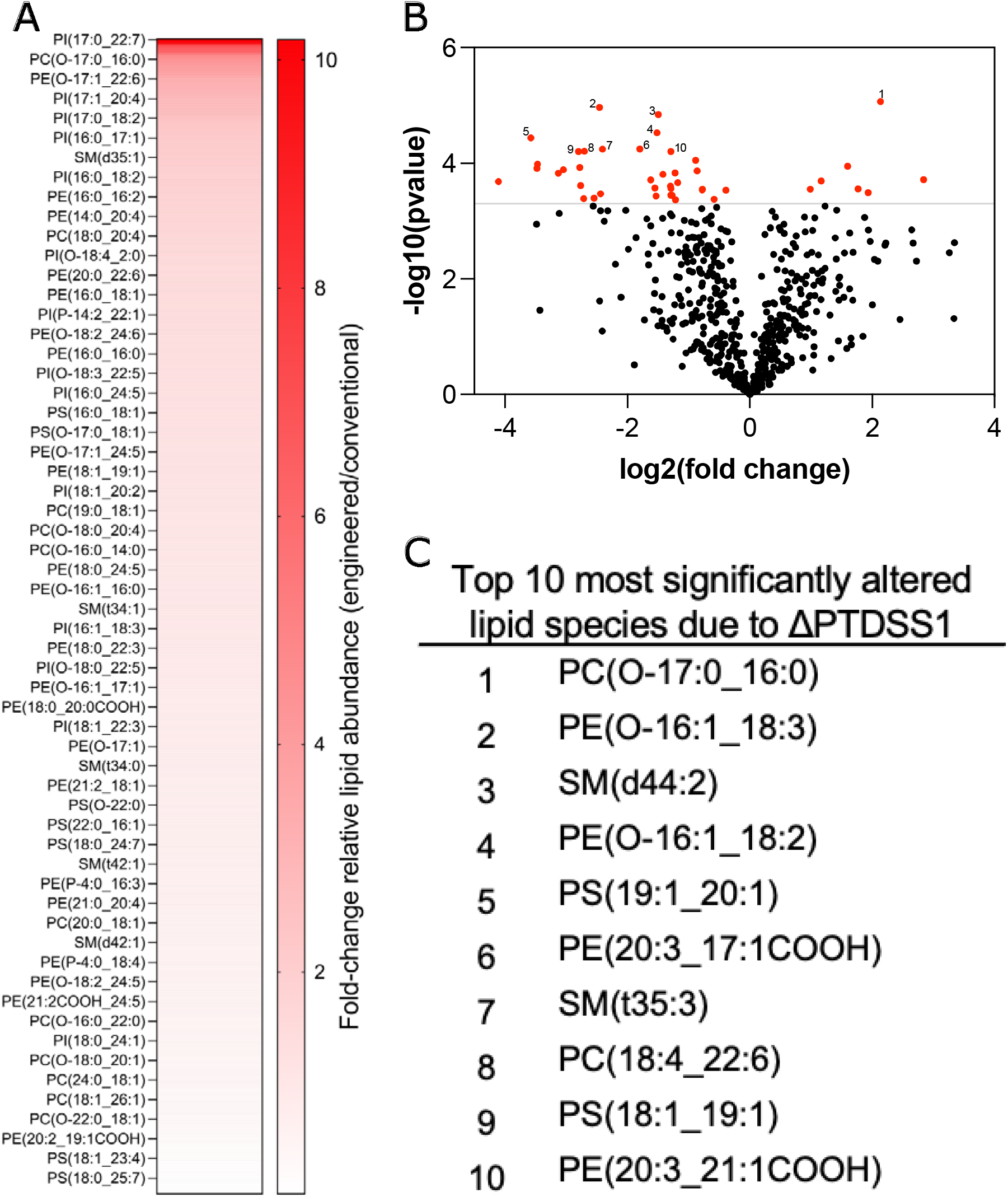
HEK293T lacking phosphatidylserine synthase showed significant alterations in lipid composition. **(A)** Heat map shows the fold-changes in the abundance of lipid species in ΔPTDSS1 cell line relative to conventional HEK293T. PC = Phosphatidylcholine, PE = Phosphatidylethanolamine, PS = Phosphatidylserine, PI = Phosphatidylinositol, SM = Sphingomyelin. Lipid species with O-represent an ether lipid. **(B)** Volcano plot shows the significant (p<0.0005) changes in lipid species when comparing ΔPTDSS1 and conventional HEK293T. **(C)** Top 10 most significantly altered lipid species depicted in the volcano plot. Lipid species were analyzed in lipid extract of 20 million cells via LC-MS negative mode (n=3).

Due to its main role in membrane fluidity and vesicle trafficking, we quantified the abundance of cholesterol in HEK293T-ΔPTDSS1 and wtHEK293T. The total cholesterol content, measured from an equal number of cells, was reduced by one-third in HEK293T-ΔPTDSS1 compared to wtHEK293T cells (**Figure 2A**). The decreased cholesterol content was accompanied by a 3-fold increase in fully saturated PS species (**Figure 2B**), possibly to maintain physicochemical properties in the cell membranes. Although we did not directly measure these properties, we measured a substantial increase in the ratio of phospholipids to cholesterol (**Figure 2C**), suggesting alterations in the bulk heterogeneity and dynamics of other membrane lipids and membrane properties. Additionally, we noted a moderate qualitative increase in the doubling time of HEK293T-ΔPTDSS1 compared to HEK293T.

**Figure 2.**
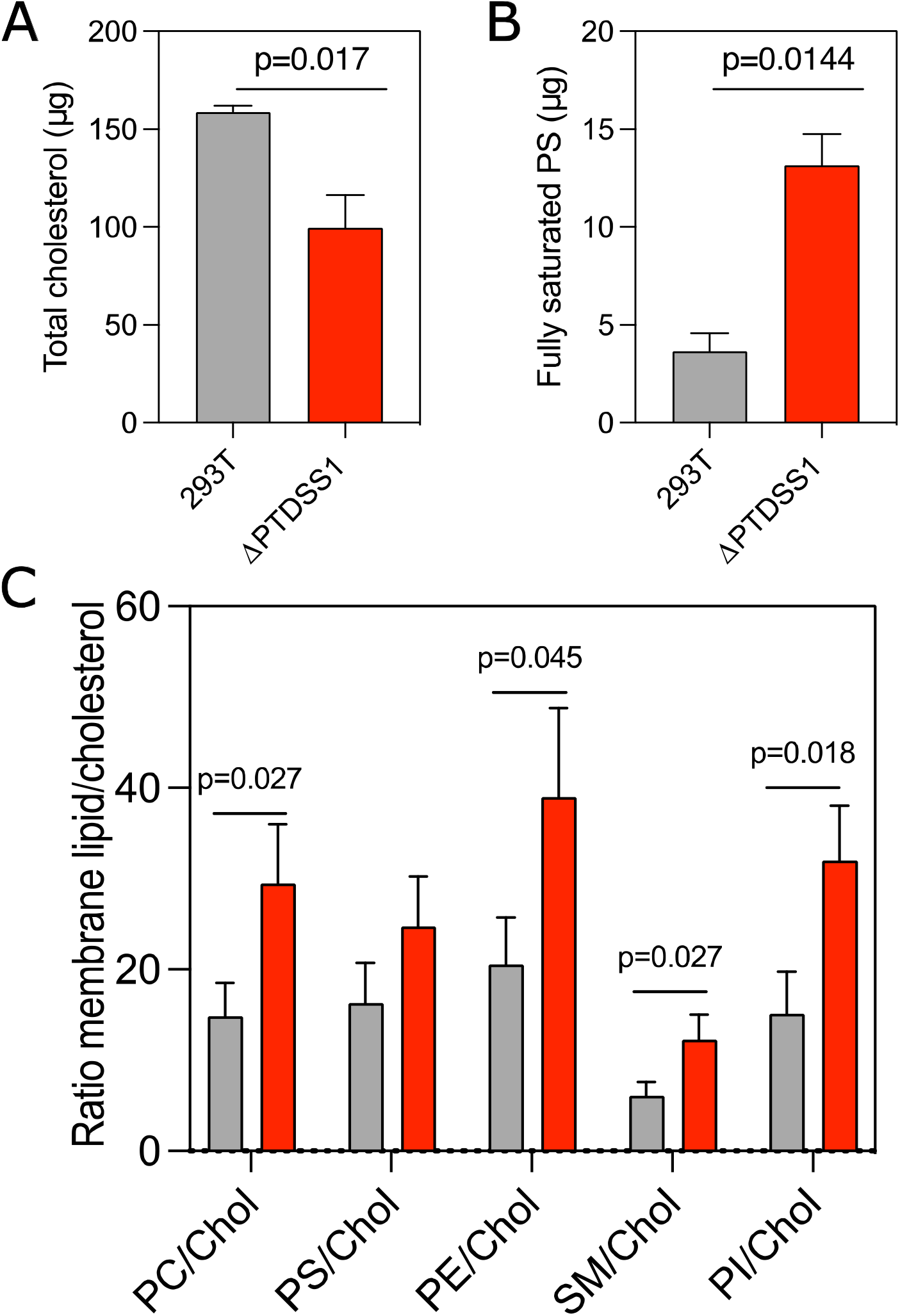
HEK29T-ΔPTDSS1 had a decrease in cholesterol abundance. **(A)** Cholesterol in the lipid extract of HEK293T-ΔPTDSS1 relative to wild type (WT) HEK293T. **(B)** The abundance of fully saturated PS species in HEK293T-ΔPTDSS1 vs WT HEK293T. **(C)** The ratio of each lipid category relative to cholesterol in HEK293T-ΔPTDSS1 (red) compared to WT HEK293T (grey).

### EV-AAV derived from PTDSS1 mutant showed enrichment of cholesterol and other membrane lipids

We characterized the lipid composition differences between the EV-AAV particles released from HEK293T and HEK293T-ΔPTDSS1. Lipidomics revealed an enrichment of several lipid species in the EV-AAV particles derived from the ΔPTDSS1 cell line (**Figure 3A**). Specifically, four lipid species had over 70-fold enrichment compared to conventional EV-AAV derived from HEK293T. Given the remarkable differences in the composition of the membrane lipids in EV-AAV derived from HEK-293T-ΔPTDSS1, we refer hereafter to these particles as mEV-AAV.

**Figure 3.**
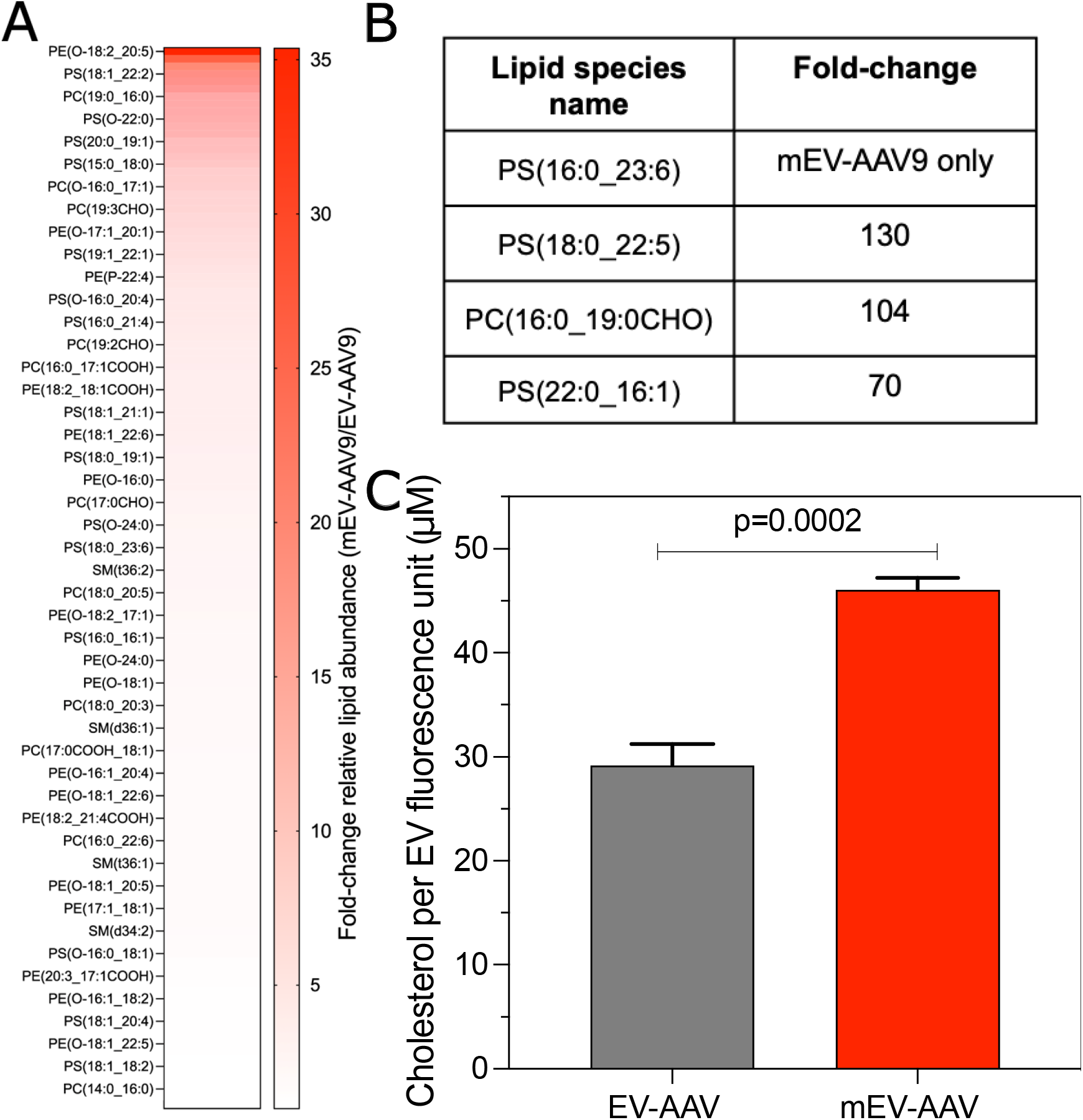
Membrane-engineered enveloped-AAV (mEV-AAV) had a different lipid composition than conventional enveloped AAV vectors (EV-AAV). **(A)** Individual lipid species relative abundance when comparing mEV-AAV and EV-AAV. **(B)** Most abundant lipid species in mEV-AAV relative to EV-AAV vectors. **(C)** EV-AAV and mEV-AAV cholesterol levels.

Among the several alterations in the relative abundance of lipid species, we detected phosphatidylserine (16:0_23:6) only in the mEV-AAV vectors. Other heterogenous PS species, with one saturated and one unsaturated fatty acid, were enriched more than 70-fold (**Figure 3B**), suggesting that the deletion of PTDSS1 produced alterations in the flux of adjacent PS biosynthesis pathways. Furthermore, we found a significant increase in the cholesterol content of mEV-AAV compared to EV-AAV (F**igure 3C**). These results showed that the lipid-modified HEK-293T-ΔPTDSS1 cell line generates membrane-engineered EV-AAV (mEV-AAV) with a unique lipid composition.

### Metabolic engineering strategy reduces free AAV in media

Due to the abundant free AAV co-released in the culture together with EV-AAV^27^, new methods for the EV-AAV manufacturing pipeline are desired. Previous studies have focused on improving downstream processes for EV-AAV purification via density gradients^23,28^. However, upstream efforts still need to be improved, specifically for cell line engineering during EV-AAV production. The lipid-modified ΔPTDSS1 cell line was tested for EV-AAV production. We produced AAV9-CBA-GFP to test the ability of ΔPTDSS1 compared to HEK293T to produce EV-AAV9 and quantify the proportion of EV-AAV9 compared to free AAV9. We employed a previously reported method of size exclusion chromatography to fractionate the supernatants and discriminate between EV-AAV9 and AAV9 fractions for either cell line^23^.

The overall yields of AAV9 vector genomes in the crude cell lysate and media were reduced by 8.2-fold and 34.2-fold, between HEK293T and HEK293T-ΔPTDSS1, respectively (**Figure 4A**). This was reflected in a 7.3-fold decrease in total yields of SEC-purified EV-AAV9 in HEK293T-ΔPTDSS1 compared to HEK293T (**Figure 4 B,C**).In agreement with our prior work for EV-AAV production, the vector genome (vg) yields of EV-AAV9 were only 25% of free AAV9 when HEK293T cells were employed (**Figure 4B**). In contrast, the metabolic engineering-guided strategy, ΔPTDSS1, shifted the ratios, resulting in an 8.2-fold larger yield of EV-AAV compared to free AAV in the culture media (**Figure 4 C,D)**. Importantly, the amount of free AAV was reduced by 300-fold in ΔPTDSS1 compared to HEK293T (**Figure 4 B,C**). Taking together, the lipid-modified ΔPTDSS1 cell line improved the purity of EV-AAV over AAV by 42.7-fold in the culture media compared to the traditionally used HEK293T with the caveat of decreased total EV-AAV by 7.3-fold.

**Figure 4.**
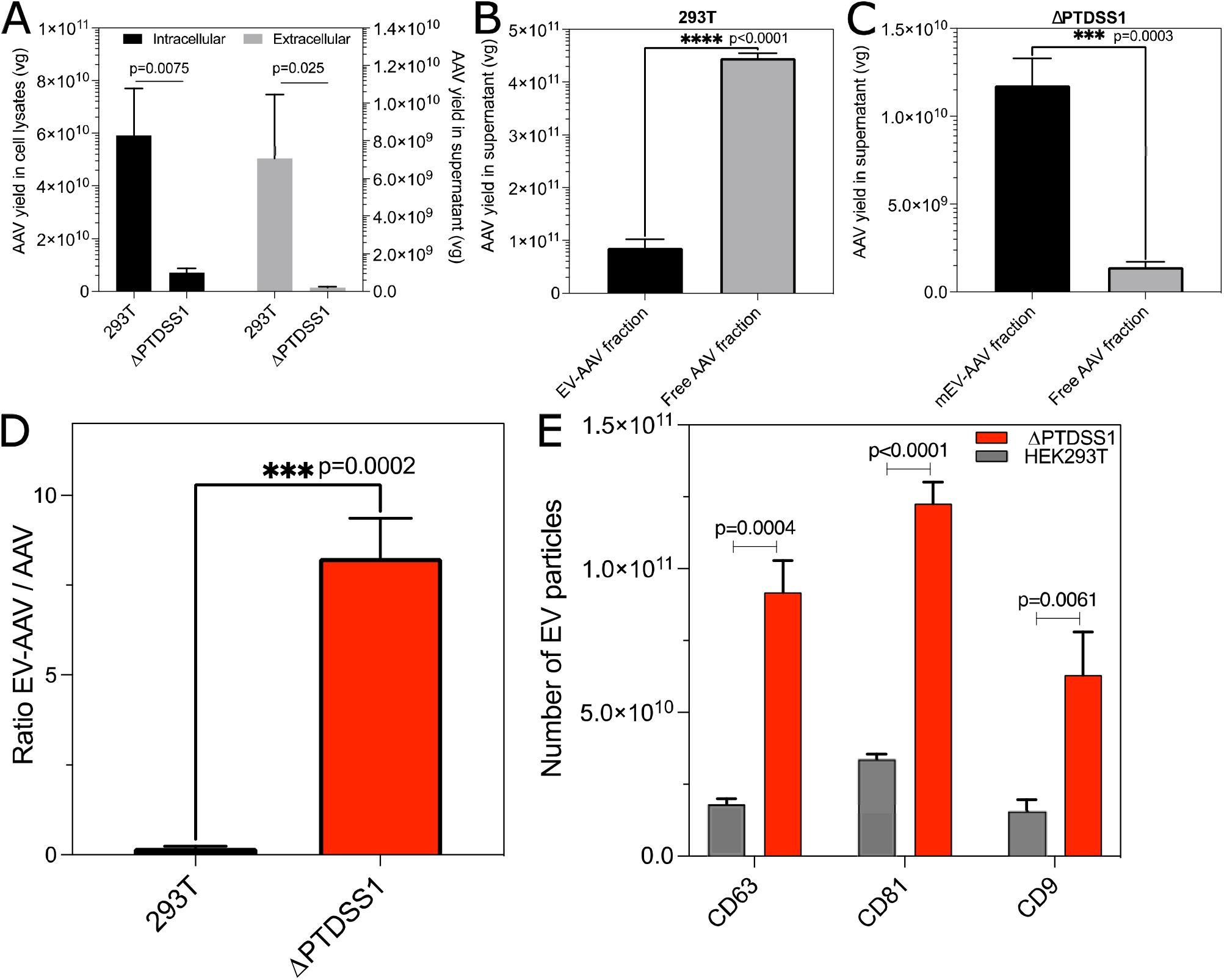
HEK293T-ΔPTDSS1 produced a higher proportion of enveloped AAV relative to free AAV. **(A)** The total AAV vg yield in the supernatant and cell lysate of HEK293T-ΔPTDSS1 compared to regular HEK293T. Size exclusion chromatography (SEC) was used to separate the fraction of EV-AAV (F1-F4) from the fractions of free AAV (F5-F12) in the supernatant of both HEK293T **(B)** and HEK293T-ΔPTDSS1 **(C)** when producing AAV. **(D)** Ratio of EV-AAV to free AAV in WT HEK293T vs HEK293T-ΔPTDSS1 **(E)** Multiplex affinity capture via Exoview was used to count extracellular vesicle (EV) markers in the fraction of EV-AAV (F1-F4) purified from HEK293T-ΔPTDSS1 (red) and the HEK293T (gray).

**Figure 5.**
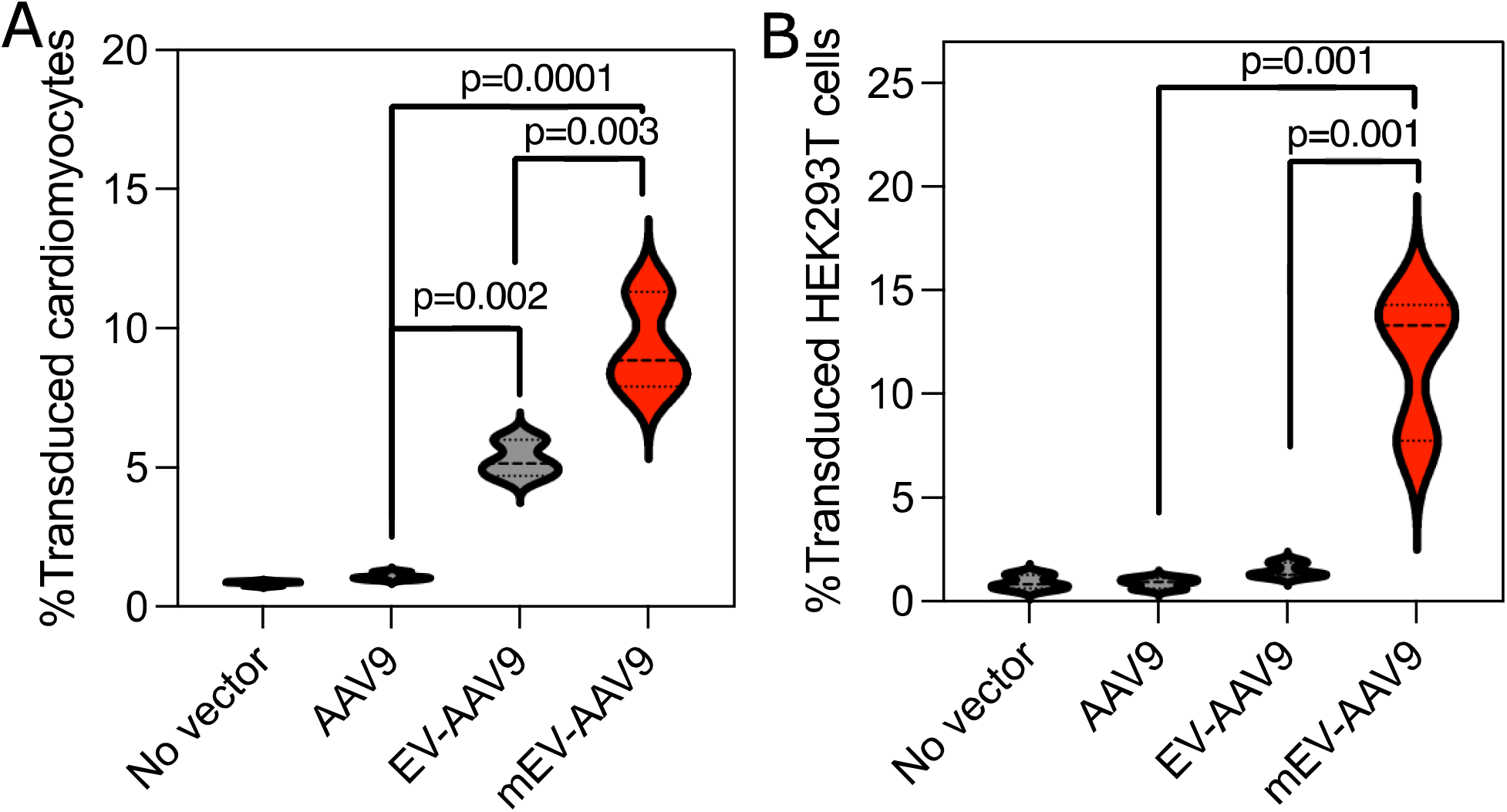
The membrane-engineered enveloped-AAV (mEV-AAV) improved transduction in human cell lines AC16 and HEK293T. The transduction of **(A)** human cardiomyocytes (AC16) or **(B)** HEK293T cells by mEV-AAV9, EV-AAV9, or AAV9 vectors encoding tdTomato. Experiments were performed using a 5.4×10^4^ vg/cell dosage with three independent replicates/ vector/cell line. Transduction (%tdTomato positive cells) was determined via flow cytometry three days after vector exposure. Statistical difference was determined after performing ANOVA and Tukey test analyses. Tukey HSD p-values are shown.

We quantified the number of EV-marker proteins in the EV-AAV fraction to determine whether the observed higher abundance of EV-AAV relative to free AAV in ΔPTDSS1 is possibly due to a higher abundance of EV. We found that the EV-AAV fraction of ΔPTDSS1 showed higher EV-marker proteins than the EV-marker protein abundance in the EV-AAV fraction of HEK293T **(Figure 4E**). These results suggest that the ability of ΔPTDSS1 to produce EV-AAV with greater purity (i.e. less free AAV) may be due to its propensity to release more EV than traditional HEK293T cells, likely owing to its engineered lipid composition.

### Membrane-engineered EV-AAV showed enhanced transduction levels in two human cell lines

The metabolic engineering strategy that enhanced EV-AAV purity during manufacturing significantly changed the lipid composition of the enveloped vectors mEV-AAV, which we reasoned may affect their interactions with cells. Consequently, we decided to test the functionality of mEV-AAV9 in two human cell lines, HEK293T cells and AC16 human cardiomyocytes. The serotype we employed as a demonstration was AAV9, and the transgene expression cassette was AAV-CAG-tdTomato. We performed a head-to-head transduction with equal doses (5×10^4^ vg/cell) of AAV9, EV-AAV9, and mEV-AAV9. Consistent with previous reports showing that EV-AAV can improve the transduction efficiency of conventional AAV in AC16 cardiomyocytes^29^ we found a 5-fold transduction improvement for cardiomyocytes when using EV-AAV compared to conventional AAV (**Figure 5A**). Engineered mEV-AAV further improved cardiac transduction by 1.8-fold compared to EV-AAV and 8.5-fold relative to traditional AAV (**Figure 5A**). The enhanced transduction ability of mEV-AAV was also measured in the HEK293T cell line. We found an over 13– and 7-fold increase in transduction due to mEV-AAV compared to traditionally used AAV and conventional EV-AAV, respectively (**Figure 5B**).

### Membrane-engineered EV-AAV showed resistance against AAV-neutralizing antibodies

A challenge for all AAV-based gene therapies is their neutralization by pre-existing anti-AAV antibodies in patients. EV-AAV vectors have shown an increased resistance against neutralizing antibodies^9–11^. To determine the resistance of mEV-AAV9 against neutralizing antibodies, we used an *in-vitro* neutralization assay^23^. We tested the ability of AAV9, EV-AAV9, and mEV-AAV9 to transduce cells in the presence of different concentrations of intravenous immunoglobulins (IVIg). Consistent with previous reports and the rationale that the viral envelope contributes to protection against the antibodies, both EV-AAV9 and mEV-AAV9 showed resistance against the highest tested concentration of IVIg relative to naked AAV (**Figure 6**). For example, at a 1:250 dilution of IVIg, conventional AAV9 displayed only 13.4% resistance to neutralization, whereas mEV-AAV9 displayed 50% resistance (**Figure 6**).

**Figure 6.**
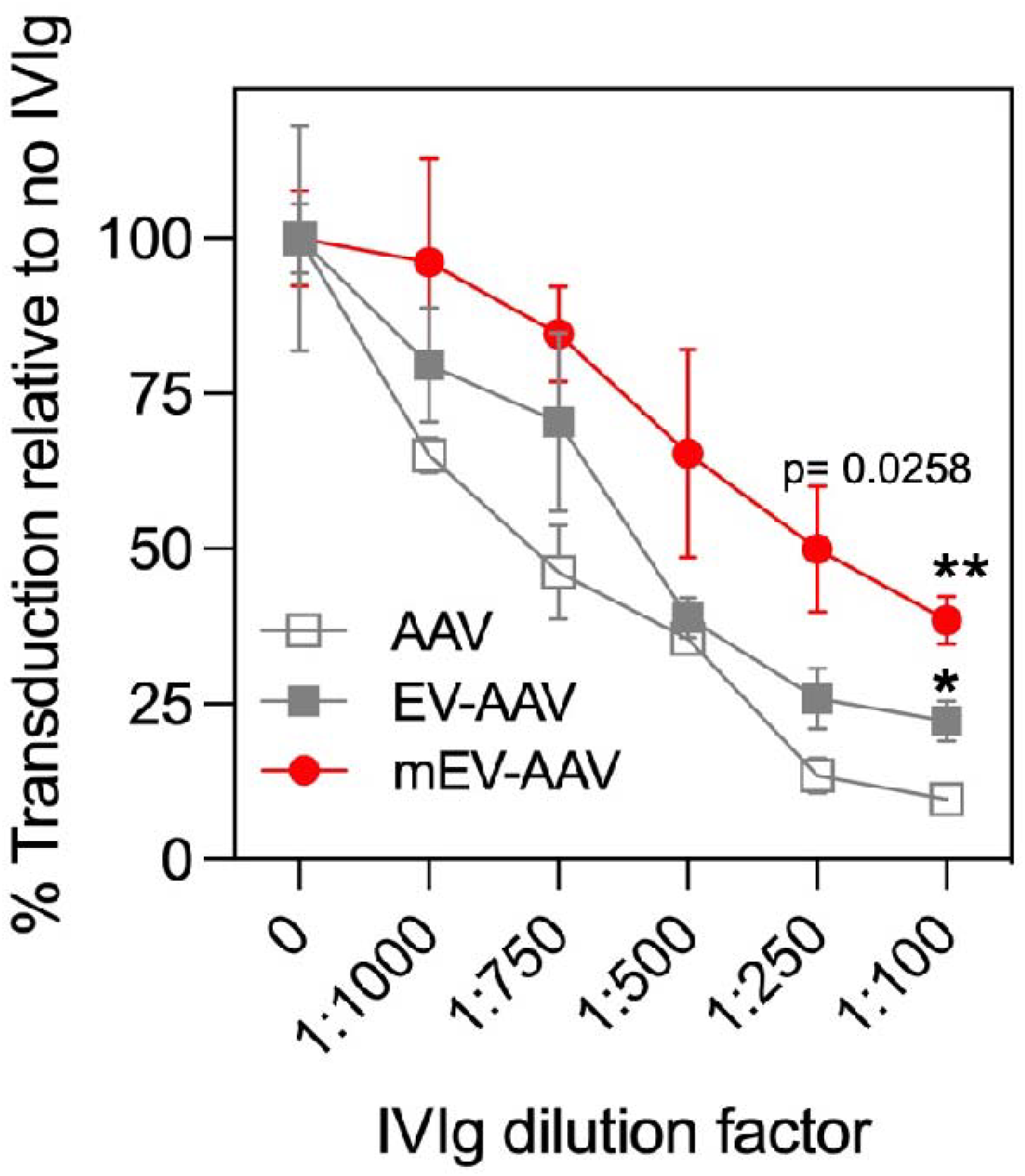
Membrane-engineered enveloped-AAV (mEV-AAV) showed resistance against neutralizing antibodies. Various concentrations of intravenous Immunoglobulins (IVIg) were incubated with AAV (empty square), EV-AAV (grey square) or mEV-AAV (red circle) before transducing HeLa cells. All vectors harbor the AAV-CBA-firefly luciferase (FLuc) transgene expression cassette. Luminescence was measured to determine transduction for each treatment. Transduction of HeLa cells with the various concentrations of IVIg was normalized to the transduction with no IVIg. The statistical difference is indicated when examining the transduction of mEV-AAV or EV-AAV relative to the AAV condition at the same concentration of IVIg. **p=0.0032, *p = 0.036. Error bars indicate the standard error of the mean (SEM). Data represents the mean of three biological replicates.

### Intracranial injection of membrane-engineered EV-AAV resulted in robust transduction in various brain regions of adult mice

We performed intraparenchymal bilateral injections into the striatum of adult C57BL/6 mice using the lipid-modified mEV-AAV9-CBA-GFP vector. Robust transduction in the brain parenchyma was observed four weeks post-injection. Transduction of cells with the morphology of neurons was observed in different brain areas, including cortical white matter, dorsal striatum (**Figure 7A**), fimbria, and internal capsule (**Figure 7B**). Furthermore, mEV-AAV9 transduction was present across various coronal sections spanning the most rostral (interaural = 5.58 mm, bregma = 1.78 mm) and most caudal (interaural = 1.00 mm, bregma = –2.80 mm) sections (**Figure 7C**).

**Figure 7.**
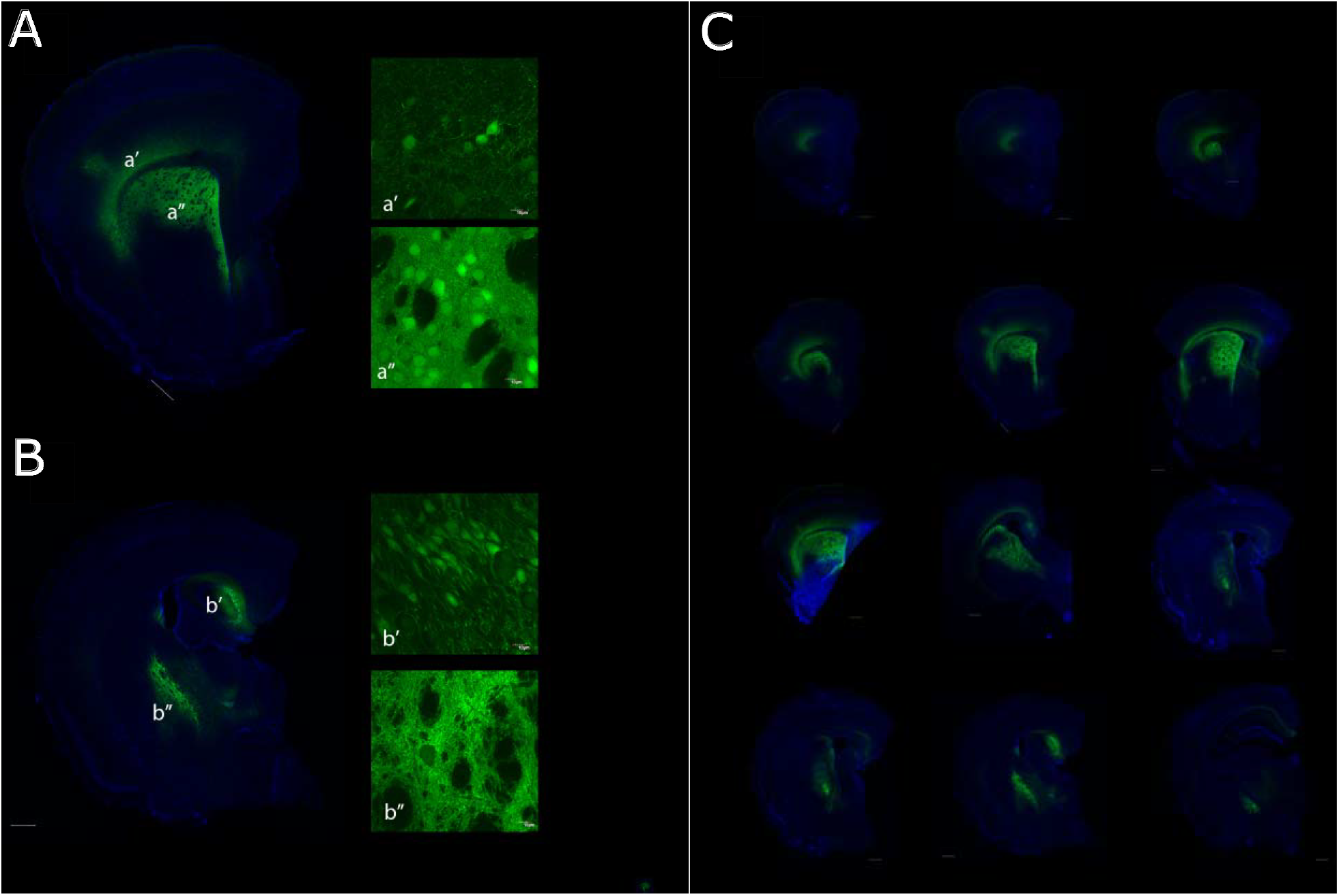
Membrane-engineered enveloped-AAV (mEV-AAV) showed robust transduction in different brain areas after intracranial injection in adult mice. **(A)** Coronal section showing cells in the cortical white matter (a’) and dorsal striatum (a’’). **(B)** Coronal section showing cells in fimbria (b’) and internal capsule (b’’). **(C)** An array of coronal sections of the brain injected mEV-AAV. All representative images show mEV-AAV9 harboring AAV-CBA-GFP transgene expression cassette. Three C57BL/6 mice were used.

## Discussion

We applied metabolic engineering concepts to disrupt the intermediate and non-essential steps for PS and PC biosynthesis, re-directing the metabolic flux towards adjacent membrane lipid biosynthesis pathways. Simultaneous targeting of PC and PS with the least possible genetic modifications was of interest as these are the most abundant lipids in the outer and inner leaflets of cellular membranes, respectively.

Lipids and proteins use the inner and outer leaflets of cellular membranes during membrane vesicle trafficking to migrate across cellular membranes of organelles and within the lipid bilayer to the opposite leaflet. Furthermore, the membrane lipid composition is driven via membrane vesicle trafficking to maintain cell homeostasis^30,31^. We anticipated that an upstream genetic strategy targeting the main components of the outer and inner leaflets could also drive alterations in membrane vesicle trafficking and, thus, in EV release and potentially in EV-AAV production.

This genetic strategy significantly increased EV production (**Figure 4E**), possibly leading to the observed higher abundance of EV-AAV compared to free AAV when HEK293T-ΔPTDSS1 was used for AAV production (**Figure 4C**). However, the increase in EVs might be due to cellular stress^32^. While we are uncertain as to the mechanism of decreased AAV production in HEK293T-ΔPTDSS1, since lipid composition is critical for cellular structure and physiology, it may not be surprising that deletion of a major driver of lipid biogenesis would affect a complex process such as AAV replication and translation of at least 9 AAV proteins, which take place in various membraned organelles.

Lipid metabolism modifying genes have been attributed to alter the size and thus the capacity of the endoplasmic reticulum (ER) to produce recombinant proteins in CHO cells^33^. The authors highlighted the role of the global transcriptional regulator, sterol regulatory element binding factor 1 (SREBF1), and the cholesterol content in ER to produce recombinant proteins, as lipid turnover in the ER is essential for optimal cellular performance and accumulation of lipids can lead to unfolded protein response^34^. Deleting the ER-resident PTDSS1 led to decreased cellular cholesterol content (**Figure 2A**), which, in line with evidence for membrane-associated accessory protein (MAAP) within the ER^35^, may have also decreased the ER capacity for AAV production. Additional studies are needed to characterize lipid-associated mechanisms for recombinant protein expression and ER functionality, such as the unfolded protein response, ER-associated chaperones, lipotoxicity, and lipid droplet formation^36^.

Despite the reduced AAV production capacity of HEK293T-ΔPTDSS1(**Figure 4A**), this cell line exhibited an increased ratio of EV-AAV to free AAV (**Figure 4C**). This improvement can contribute to simplifying downstream unit operations for commercial-scale EV-AAV production. Specifically, if less free AAV is present in the media, fewer steps will be required for their depletion, improving process time and yield. Furthermore, the resultant mEV-AAV had a unique lipid composition characterized by a higher abundance of cholesterol (**Figure 3A-C**). The decrease of cholesterol in HEK293T-ΔPTDSS1 and the increase of cholesterol content in the mEV-AAV, strongly suggest that the membrane lipid composition of the envelope surrounding AAV can be modulated. Furthermore, as mentioned earlier, the role of cholesterol in cellular membranes for protein expression and folding could be used as a strategy for envelope engineering of EV-AAV. Here, we decided to characterize the gene delivery functionality of this novel composition of mEV-AAV compared to EV-AAV. In agreement with our data, previous studies have shown EV-AAV transduction of cardiomyocytes both in-vitro and *in-vivo*^14^. The further improvement in mEV-AAV transduction of hAC16 and HEK293T cells might be due to the unique envelope composition. However, a potential decrease in the abundance of AAV particles non-covalently bound to the outer surface of the EV in mEV-AAV vectors might be another possibility for the observed improvements in gene delivery (**Figure 5**) and resistance against antibody neutralization (**Figure 6**) compared to conventional EV-AAV. Further studies on the composition and different purification methods will be relevant for elucidating the most critical parameters for improving gene delivery and resistance against neutralizing antibodies in the EV-AAV platform. Remarkably, enveloped-AAV (EV-AAV and mEV-AAV) outperformed conventional AAV for gene delivery in cultured cells (**Figure 5**), resistance against neutralizing antibodies (**Figure 6**), and also showed robust *in-vivo* transduction after local administration in the murine brain (**Figure 7**). Further head-to-head comparisons of versions of EV-AAV, LNP ^4,37^, retrovirus-like particles^37^, and other gene delivery systems will be essential to develop a safe vector with the most clinically relevant properties. We anticipate that further upstream efforts, including additional metabolic engineering strategies, can positively impact the large-scale production of clinically relevant gene therapy vectors.

## Conclusion

This study demonstrates that metabolic engineering can be applied to improve EV-AAV vector development. Specifically, HEK293T-ΔPTDSS1 produced a higher proportion of EV-AAV relative to free AAV by 42.7-fold while decreasing the amount of free AAV by 300-fold in the supernatant. The membrane-engineered enveloped vectors (mEV-AAV) had a unique lipid composition, including increased cholesterol content and higher transduction efficiency in human cardiomyocytes AC16 compared to conventional EV-AAV and regular non-enveloped AAV. Additionally, mEV-AAV vectors showed beneficial properties of enveloped-AAV (EV-AAV), such as increased resistance to anti-AAV antibodies and robust transduction *in-vivo* after local intracranial administration in mice. These findings strongly suggest that metabolic engineering can be used to address existing limitations in EV-AAV-based gene therapies.

## Acknowledgments

This work was partially supported by the American Society of Gene and Cell Therapy (ASGCT) Underrepresented Population Fellowship Award (to M.C.S.), and by the NIH R01 grant NIH [DC017117] (to C.A.M.). We thank Jennifer X. Wang at the Harvard Center for Mass Spectrometry for her assistance in running and analyzing the lipidomic analyses.

## Author contributions

M.C.S and C.A.M. conceived of the study and wrote the manuscript. P.E. M.C., C.N., D.D.L.C, E.W., D.M.Mc.C, and M.C.S performed experiments and analyzed data. P.G.B. and C.A.M. provided critical insight into experimental design and analysis of the data.

